# Event-Related Warping: A Toolbox for Temporal Alignment and Jitter Correction in Sequential Experimental Paradigms

**DOI:** 10.64898/2026.03.24.713943

**Authors:** Andrew D Levy, Peter Zeidman, Karl Friston

## Abstract

Sequential experimental paradigms are fundamental to cognitive neuroscience, yet standard event-related response analysis struggles with the temporal variability inherent to these designs. Conventional epoching treats each event within a sequence as an independent response, discarding the temporal dependencies between successive events and obscuring systematic changes in neural state that accumulate across the sequence. In order to generate responses that capture the entire sequence, it necessitates alignment across trials to correct for the inherent temporal jitter that would otherwise blur averaged responses and obscure the true sequential dynamics. Existing temporal alignment methods warp observed signals directly, making them vulnerable to correlated noise and potentially disrupting multichannel temporal relationships essential for connectivity and causal analyses. Event-Related Warping (ERW) addresses these limitations by aligning template functions encoding experimental event structure rather than neural signals themselves. Templates constructed from event onsets and durations undergo smooth monotonic warping via gradient-based optimisation, then estimated trajectories are applied uniformly across all channels, preserving inter-channel timing relationships and causal structure. This design-level alignment exploits experimentally observable jitter whilst maintaining signal integrity. Simulations with known ground truth incorporating Gaussian jitter, skewed latencies, amplitude-latency coupling, and multi-parameter dependencies yielded standardised root-mean-square errors (sRMSE) of 0.27-0.38. Distance-weighted averaging, emphasising temporally consistent trials, provided 5-13% improvement when jitter exceeded 100 ms, with maximal benefit (≈13% reduction) under quadratic amplitude-latency coupling. Empirical validation using an auditory go/no-go dataset with cue-to-target intervals of 1.5-4.1 seconds demonstrated that ERW recovers jittered target-locked responses with comparable fidelity (sRMSE 0.24-0.51) to conventional epoching of time-locked events, whilst preserving inter-channel lag relationships (cross-covariance sRMSE 0.63-0.82). ERW thus extends standard trial averaging to scenarios where temporal variability would otherwise preclude coherent response recovery, supporting investigation of temporally extended processing in ecologically valid paradigms whilst maintaining compatibility with established ERP frameworks and downstream connectivity analyses.

## 1 Introduction

Sequential paradigms form a cornerstone of experimental psychology and cognitive neuroscience, encompassing designs from simple serial reaction time tasks (Robertson 2007) to complex sequential decision-making protocols (Roijers et al. 2013). Whether examining attentional dynamics through rapid serial visual presentation (Klee, Memmott, and Oken 2024), probing cognitive control via task-switching (Davidson et al. 2006), or investigating predictive processing through oddball sequences (Garrido et al. 2008), researchers routinely employ experimental designs in which multiple stimuli are presented in temporal succession. These designs are essential for capturing the inherently dynamic nature of cognitive processes, from perceptual integration (Lisi, Morgan, and Solomon 2020) and priming effects (Nieuwenhuis et al. 2006) to evidence accumulation during decision-making (Gluth, Rieskamp, and Buchel 2012) and sustained attention monitoring (Jia et al. 2017). Neural responses to any given stimulus are fundamentally shaped by preceding context through multiple neurophysiological mechanisms: adaptation and refractoriness alter neuronal excitability (Sawamura, Orban, and Vogels 2006); neuromodulatory tone shifts neural gain and signal-to-noise characteristics (Auksztulewicz et al. 2017); ongoing oscillatory states determine the phase and amplitude context for new inputs (Lakatos et al. 2005); neurotransmitter dynamics influence synaptic efficacy (Alabi and Tsien 2012); and cumulative processes, from attention-related baseline shifts (Thut et al. 2006) to working memory maintenance (Miyake and Shah 1999) and expectation-driven preparatory states (Eimer, Velzen, and Driver 2002), create history-dependent response profiles that evolve continuously across event sequences (Friston 2000a, 2000b, 2000c).

Despite this recognition, standard EEG and MEG analysis approaches sequential experimental designs by partitioning the sequences into time-locked epochs composed of a single event type of a sequence. Whilst this epoching approach has proven successful in characterising stereotyped post-stimulus responses, it is imposed by practical necessity: temporal variability inherent to sequential paradigms, including varying inter-stimulus intervals and reaction-time jitter, precludes simple continuous modelling. By time-locking to discrete events and averaging across epochs, traditional analysis treats each stimulus as initiating a relatively independent neural response, maximising signal-to-noise ratio for post-stimulus components but discarding the tonic shifts, sustained preparatory states, and history-dependent dynamics that evolve continuously without resetting at event boundaries. The ongoing debate surrounding baseline correction exemplifies this tension, reflecting implicit assumptions about whether pre-stimulus periods represent neutral reference states or meaningful task-relevant dynamics (Alday 2019; Tanner et al. 2016; Maess, Schröger, and Widmann 2016).

Methodological advances now enable analysis of extended timescales: dynamic causal modelling decomposes fast and slow neural dynamics (Medrano, Friston, and Zeidman 2024), hierarchical Bayesian frameworks accommodate temporally structured priors (Kuzin, Isupova, and Mihaylova 2018), and convolution-based regression approaches model overlapping responses (Litvak et al. 2013). Simultaneously, neuroimaging increasingly employs ecologically valid naturalistic paradigms with continuous sensory stimulation or self-paced behaviour (Virk, Letendre, and Pathman 2024; Vanderwal, Eilbott, and Castellanos 2019). Modern annotation tools enable precise event coding (Su, Hairston, and Robbins 2018; Diachenko et al. 2022). Capitalising on these developments requires alignment methods that accommodate experimental jitter whilst recovering high signal-to-noise trial averages suitable for sophisticated downstream analyses.

Temporal alignment techniques have been developed to address trial-to-trial variability in electrophysiological recordings. Dynamic time warping (DTW) algorithms employ dynamic programming to align time series by minimising point-wise distances, widely applied to EEG for clustering waveforms and discriminating latency differences (Huang and Jansen 1985; Zoumpoulaki et al. 2015). Cross-correlation methods, such as the Woody filter, iteratively estimate single-trial latencies to sharpen jittered components (Woody 1967; Kutas, McCarthy, and Donchin 1977). More recently, diffeomorphic warping methods have been introduced to achieve smooth, invertible transformations (Martínez 2003; Rivet and Cohen 2016). These approaches share a common strategy: they warp the observed neural signals to achieve temporal alignment. Whilst this can successfully reduce jitter and improve signal-to-noise ratio, signal-warping methods are vulnerable to correlated noise and distortion of multichannel temporal structure (Lysiak 2021). Critically, these methods are data-driven rather than informed by experimental event timing, aligning based on signal features rather than the known temporal variability inherent to the paradigm design.

Alternative approaches address the distinct problem of overlapping event-related responses. Decomposition methods such as Residue Iteration Decomposition (RIDE) separate stimulus-locked and response-locked components through iterative realignment and subtraction (Ouyang et al. 2011), whilst regression-based convolution methods model overlapping responses (Litvak et al. 2013). These techniques successfully disentangle mixed contributions from temporally proximate events but yield independent estimates for each component rather than unified sequential responses that preserve the continuous temporal cascade of sensorimotor processing.

We introduce Event-Related Warping (ERW) to address these limitations (poor handling of multichannel data, sensitivity to noise, and failure to exploit known event structure) by performing temporal alignment on latent event representations rather than observed signals. ERW constructs template functions encoding event onsets and durations from the experimental design, then estimates smooth warping trajectories that align these templates across trials. Critically, the jitter being corrected is experimentally observable: variable reaction times, stimulus-onset asynchronies, or behavioural response latencies. ERW does not attempt to recover unobservable neurophysiological latencies or endogenous sources of variability, which remain the domain of data-driven methods. Once estimated from template alignment, warping trajectories are applied uniformly to all simultaneously recorded channels, preserving inter-channel temporal relationships essential for connectivity and causality analyses. Standard trial averaging then reconstructs unified sequential responses spanning multiple events, compatible with standard ERP interpretation, whilst accommodating substantial experimental jitter.

We establish ERW’s validity through complementary analyses. Simulation studies with known ground truth assess recovery accuracy under systematic temporal variability. Empirical validation uses an open-access auditory go/no-go EEG dataset to test whether ERW recovers sequential potentials whilst preserving inter-channel timing. Together, these establish ERW as a principled method for paradigms where sequential events exhibit substantial experimentally induced variability, enabling investigation of temporally extended processing whilst maintaining compatibility with established ERP frameworks.

## 2 Methods

### 2.1 Timeseries Alignment

#### 2.1.1 Background Methods

Given two sequences, Dynamic Time Warping (DTW) computes an optimal alignment path that minimises the cumulative distance between them (*x* and *x*′):

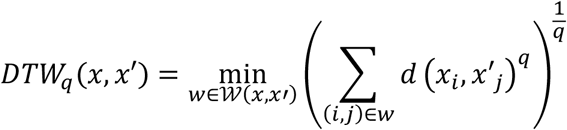

where *d*(·,·) is a distance measure (e.g., Euclidean when *q* = 2), and *w* = ((*i*_1_, *j*_1_), …, (*i*_*L*_, *j*_*L*_)) is a warping path of length *L* within the admissible set 𝒲(*x, x*′), subject to monotonicity, continuity and boundary constraints.

Extending DTW to *N* sequences introduces an averaging problem: find a centroid sequence ŷ that minimises total DTW distance:

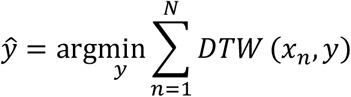

DTW Barycentre Averaging (DBA) addresses this via alternating align-and-average steps (Petitjean, Ketterlin, and Gançarski 2011). Each iteration aligns all sequences to the current average using standard DTW, then updates the average by aggregating aligned points. Whilst principled, DBA is sensitive to initialisation and often converges on weak local minima due to DTW’s discrete, non-differentiable nature.

Generalised Time Warping (GTW) reformulates DTW averaging as a joint optimisation problem by representing warping paths as weighted combinations of monotonic basis functions, enabling gradient-based optimisation (Zhou and De la Torre 2012). However, GTW still relies on discrete sampling, limiting alignment smoothness and precision.

#### 2.1.2 Trainable Time Warping

Trainable Time Warping (TTW) extends GTW by making alignment fully differentiable, enabling direct gradient-based optimisation of temporal parameters (Khorram, McInnis, and Mower Provost 2019). Unlike GTW’s discrete signals, TTW constructs continuous-time approximations using differentiable interpolation kernels. Smooth, time-dependent warping functions parameterised via truncated Discrete Sine Transform (DST) allow non-linear alignment whilst constraining the solution space to reduce overfitting. Warped signals are resampled onto a common axis and averaged pointwise. This formulation jointly optimises the entire pipeline (interpolation, warping and averaging) using standard gradient methods. By avoiding discrete mappings, TTW achieves superior alignment fidelity and smoother solutions than GTW, particularly for complex or temporally non-uniform distortions.

##### 2.1.2.1 Technical details

In this work, we implemented a parabolic interpolation kernel *ϕ*(*t*) for continuous-time reconstruction, offering computational efficiency over the sinc kernel used in the original TTW procedure (Khorram, McInnis, and Mower Provost 2019). This is enabled by the sparse nature of the event representations aligned in this procedure (see section Event-Related Warping)

The parabolic kernel comprises three overlapping quadratics with support on [−1.5,1.5]:

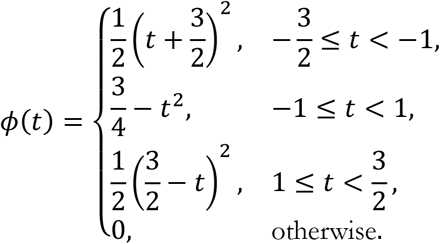

Let *x*_*n*_[*t*] denote a discrete signal of *N* trials sampled at interval *T*_*s*_. Its continuous-time approximation 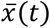 is:

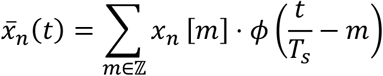

A fixed temporal shift *τ*_*n*_ ∈ ℝ can be incorporated:

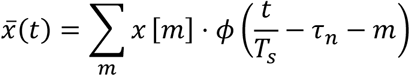

Returning to discrete time by setting *t* = *tT*_*s*_ yields:

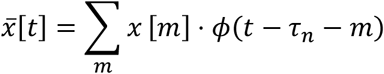

For non-linear alignment, we replace the fixed shift with a time-dependent warping function *τ*_*n*_[*t*], producing a warped signal 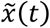:

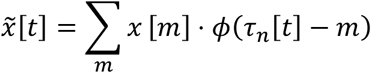

The warping function uses a DST parametrisation of order *K*:

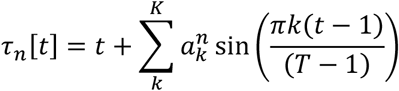

Here 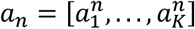 are optimised DST coefficients. We denote the full warping set as 𝒯 = {*τ*_1_, … *τ*_*N*_} with optimisation problem:

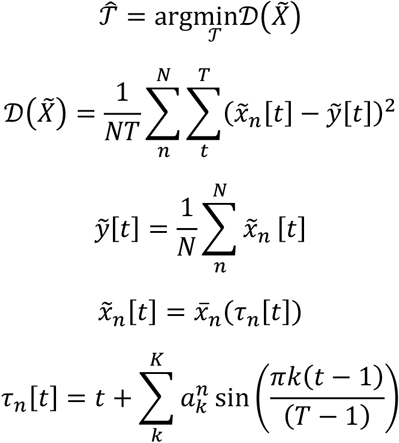

Here 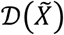 denotes the within-group mean squared error (MSE) of synchronised signals and 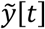 represents the sample-wise average of aligned signals.

Standard DTW constraints regularise the solution:

- **Continuity**: Enforced by limiting DST coefficients *K*. Larger *K* permits greater warping range but increases discontinuity and overfitting risk.
- **Monotonicity**: Enforced post-hoc by replacing violations where *τ*_*n*_[*t*] < *τ*_*n*_[*t* − 1] with *τ*_*n*_[*t* − 1].
- **Boundary Conditions**: Endpoints fixed at *τ*_*n*_[1] = 1 and *τ*_*n*_[*T*] = *T*, ensuring complete signal domain coverage.

Optimisation employs BFGS quasi-Newton method via MATLAB’s *fminunc*, supporting gradient-based formulation. We derive closed-form analytic gradients of 𝒟 with respect to DST coefficients *a*:

Let 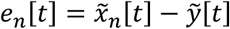. Then:

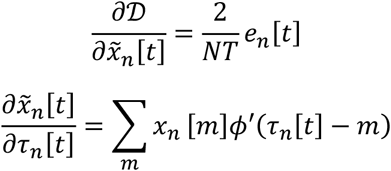

Chaining:

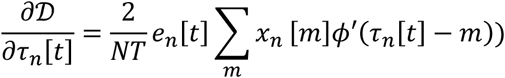

Since 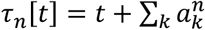 sin(*πk*(*t* − 1)/*T* − 1):

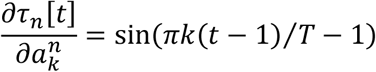

Therefore:

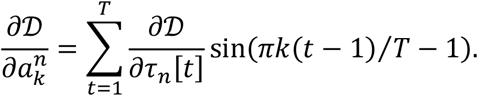

#### 2.1.3 Event-Related Warping

Whilst TTW successfully aligns timeseries by minimising point-wise differences, neuroimaging applications face additional challenges. Raw neural signals contains spatially and temporally correlated noise that can distort alignment, and multichannel recordings require consistent temporal transforms across sensors to preserve causal relationships. We therefore extend TTW to Event-Related Warping (ERW), which aligns structured event templates rather than raw signals.

A template, analogous to a design matrix, encodes the onset and duration of events within each trial. Aligning these structured representations offers several advantages: it eliminates noise influence on alignment estimation, enables uniform application of warping trajectories across all simultaneously recorded channels and modalities, and preserves underlying causal relationships between signals.

The procedure comprises four stages: template construction, template warping via TTW, application of resulting trajectories to raw data, and weighted averaging based on warping distance.

##### 2.1.3.1 Template Construction

We encode each trial’s event structure as a continuous “bump” signal. For trial *i*, we place a Gaussian bump *f*(*t, θ*) at each event onset *θ*_*ij*_, with dispersion set by event duration. Summing over *M* events yields the trial-specific template:

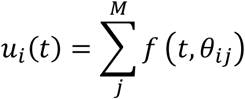

To obtain a common alignment reference, we compute the central tendency 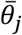 of the *j*th event’s parameters across all *N* trials (e.g., mean or median), then construct the target template ū :

Direct alignment to ū (*t*) risks local minima. We therefore generate intermediate templates that gradually transition from the original to the target timing. At step *s*, we blend parameters:

forming intermediate templates:

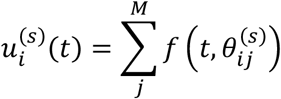

This sequence (**Figure 1. 1A**) provides a smooth optimisation path from raw (*s* = 0) to final (*s* = *S*) timing, improving convergence.

**Figure 1:**
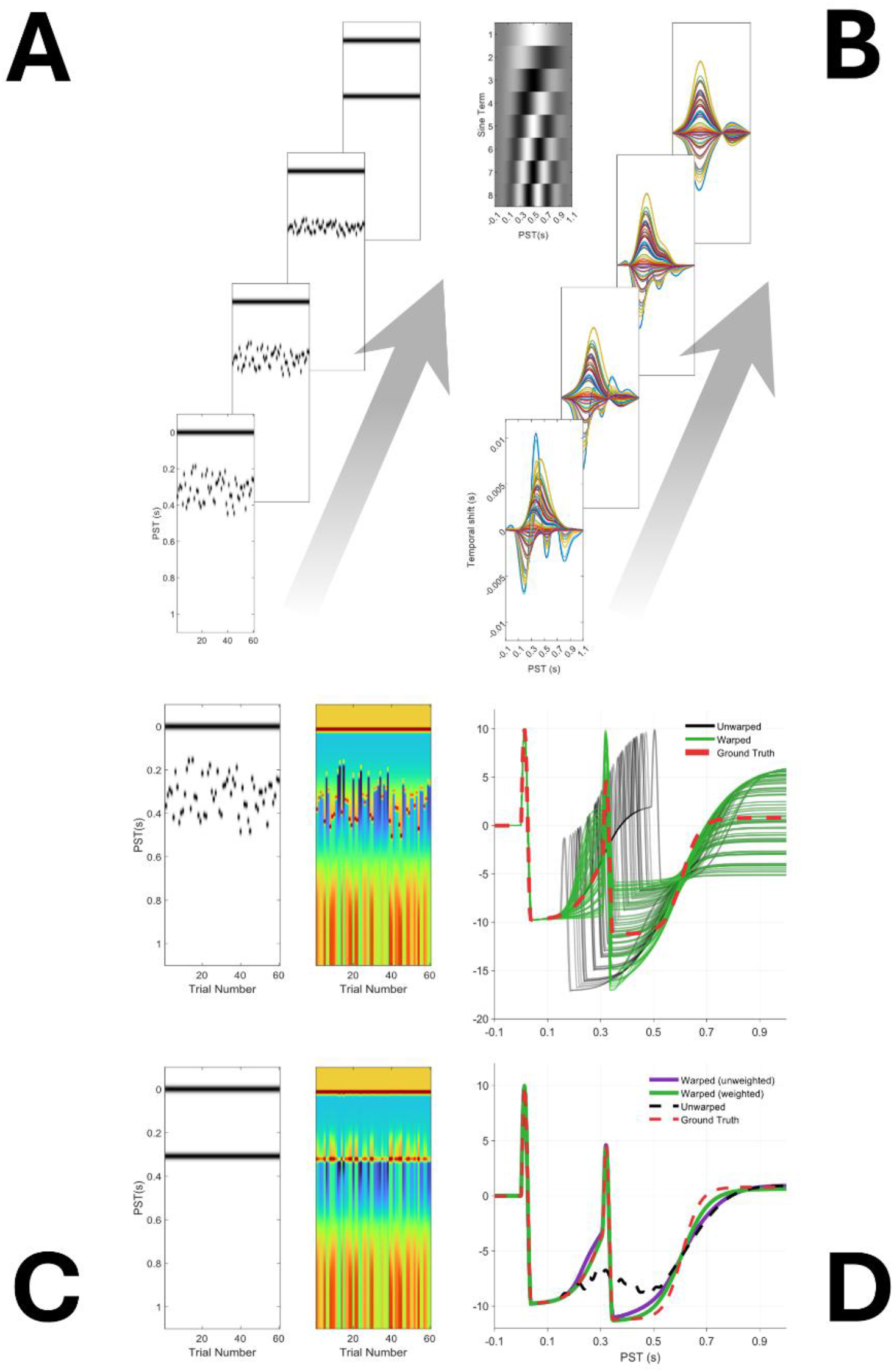
Event-Related Warping Pipeline: The four panels summarise transformation of heterogeneous trials into temporally aligned, trial-averaged responses. A. Template construction. Each raster plot depicts an intermediate event template generated by summing Gaussian “bumps” placed at annotated event onsets for each trial. From bottom to top, event parameters are linearly interpolated, gradually shifting templates from original to shared target timing. B. Warp-trajectory estimation. For every template level, a smooth, monotonic warping curve maps the current template to the next. Grey inset displays the order-8 DST basis representing these curves. Coloured traces show resulting displacement trajectories at successive refinement steps: positive values delay samples, negative values advance them. C. Signal warping. Final warp trajectories are applied to raw data. Clockwise from upper-left: (i) first-step template, (ii) unwarped trials, (iii) warped trials—now vertically aligned—and (iv) final target template. D. Trial averaging. Upper subplot—warped trials (green) overlaid on unwarped data (black) with ground-truth response (red dashed), highlighting tighter synchrony achieved by ERW. Lower subplot—conventional mean (purple) contrasted with distance-weighted mean (green), showing how down-weighting trials requiring large temporal corrections yields an average more closely following ground truth.

##### 2.1.3.2 Warp-Trajectory Estimation

At each step *s*, we align the intermediate template set *U*^(*s*)^ using TTW. Each discrete template is lifted to continuous time via the parabolic kernel, then smooth monotonic warps 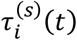 are parameterised in an order-*K* DST basis. Optimal warps are found by gradient-based minimisation of within-group MSE:

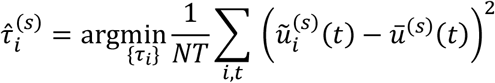

These trajectories (**Figure 1. 1B**), estimated at each level, capture the smooth, monotonic displacement aligning each trial’s template to the target at step *s*.

##### 2.1.3.3 Signal Alignment

Once per-trial warp trajectories *τ*(*t*) are estimated, we apply them directly to raw traces (**Figure 1.1C**):

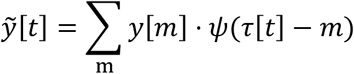

We employ modified Akima (Makimi) piecewise cubic interpolation (*ψ*) rather than the parabolic kernel. Makima preserves sharp EEG transients whilst avoiding overshoot artefacts of standard cubic splines and ensuring continuous first derivatives, maintaining event-related response morphology during resampling.

##### 2.1.3.4 Trial Averaging

After warping, we compute representative responses by averaging aligned trials (**Figure 1.1 D**). Conventional averaging treats all trials equally. Alternatively, we can weight trials by their warping distance (a scalar quantifying temporal deviation from the target) to emphasise temporally consistent trials.

We implement distance-weighted averaging as follows. For trial *n*, compute total squared temporal shift:

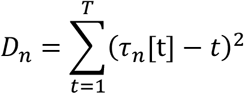

Let *σ* denote the standard deviation of {*w*_*n*_} across trials and *κ* a user-defined sharpness parameter. Then:

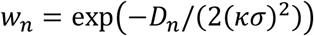

Normalise *w*_*t*_ to unity sum and compute weighted mean:

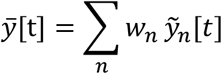

Smaller *κ* emphasises trials closely following the template, sharpening features in high-SNR data. Larger *κ* prevents noise amplification in low-SNR scenarios. We explore how *κ* interacts with SNR to optimise weighted averaging.

### 2.2 Validation via Simulation

To assess ERW’s accuracy under conditions where true temporal structure is precisely known, we conducted ground-truth validation using synthetic event-related responses. This approach provides face validity, direct quantification of whether ERW retrieves known signals after realistic perturbations. We generated canonical two-event waveforms and systematically introduced four classes of trial-to-trial variability simulating neurophysiological processes: (A) Gaussian-distributed onset jitter, (B) skewed latency distributions resembling reaction-time data, (C) amplitude-latency coupling reflecting intensity-dependent neural responses, and (D) combined multi-component coupling. By quantifying ERW’s ability to recover the original unperturbed signal across these scenarios, we establish both recovery validity (accurate parameter retrieval) and convergent validity (generalisation across multiple manifestations of temporal variability).

Each synthetic trial comprised two sequential transients: a brief oscillatory burst followed by slower recovery. These were modelled as windowed sinusoids and sigmoid functions:

We defined a canonical waveform (**Figure 2**) with two events separated by 300 ms. The sinusoidal burst had a frequency *f* = 20 Hz, duration 37.5 ms, and amplitudes 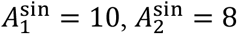. The sigmoid recovery had a slope *k* = 30, midpoint 300 ms, and amplitude 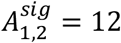. All waveforms were sampled at 500 Hz over −100 to 1000 ms. This canonical response served as basis for simulations introducing systematic trial-to-trial variability: onset jitter, amplitude scaling, skewed latency distributions and nonlinear coupling between components, testing ERW’s robustness under realistic neural dynamics.

**Figure 2:**
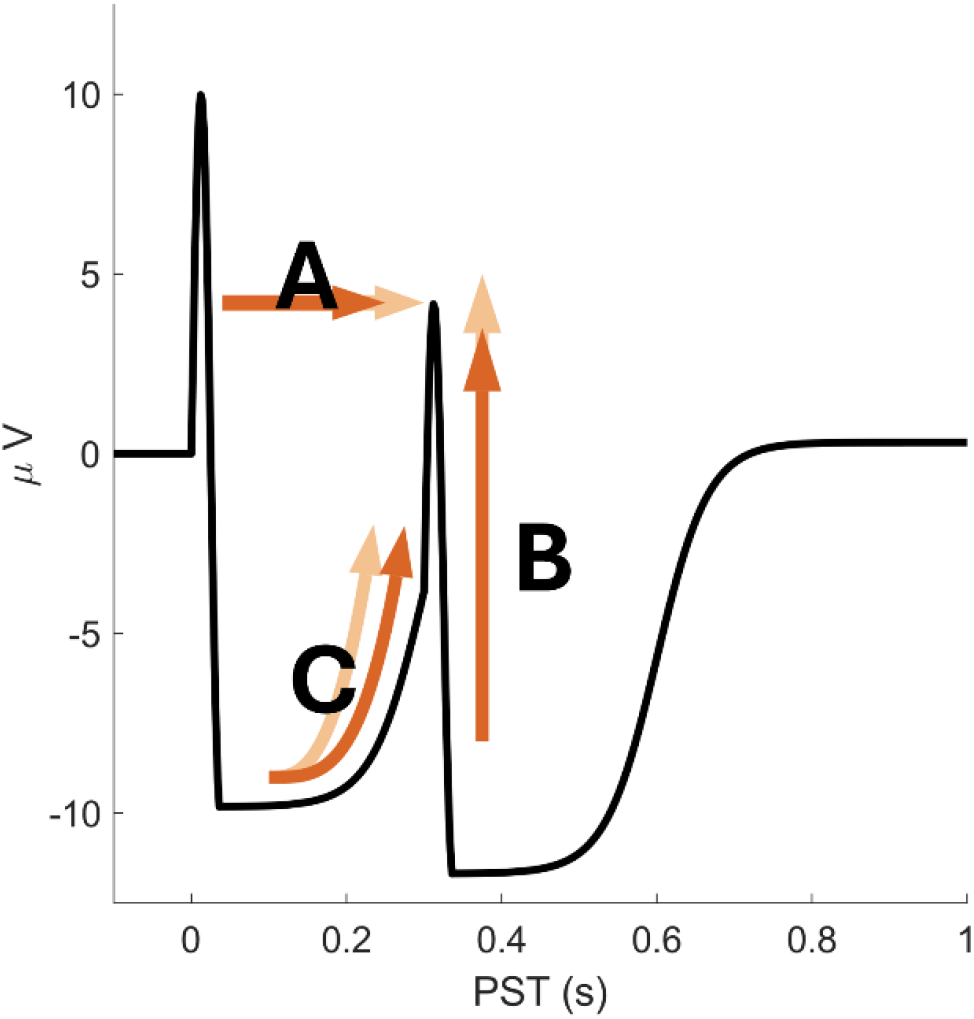
Canonical synthetic event-related waveform and variability manipulations: Black trace shows the canonical response used to validate ERW: two transient events separated by 300 ms, each comprising a brief oscillatory burst followed by slower sigmoid-shaped recovery. First burst peaks at 10 µV, second at 8 µV, both lasting 37.5 ms at 20 Hz, with recovery phases rising to 12 µV at midpoint 300 ms after each burst. Sampling at 500 Hz over −100 to 1000 ms window. Overlaid arrows indicate trial-to-trial variability types introduced: A onset latency shifts of second event, B amplitude scaling of second burst, C changes in first event’s recovery slope.

### 2.2.1 Simulation Variations

#### A. Normally distributed onset variability (Figure 3A)

To examine sensitivity to random timing jitter, we sampled second event onset latency from Gaussian distributions with standard deviations 25 ms, 75 ms and 125 ms, testing three variability levels.

**Figure 3:**
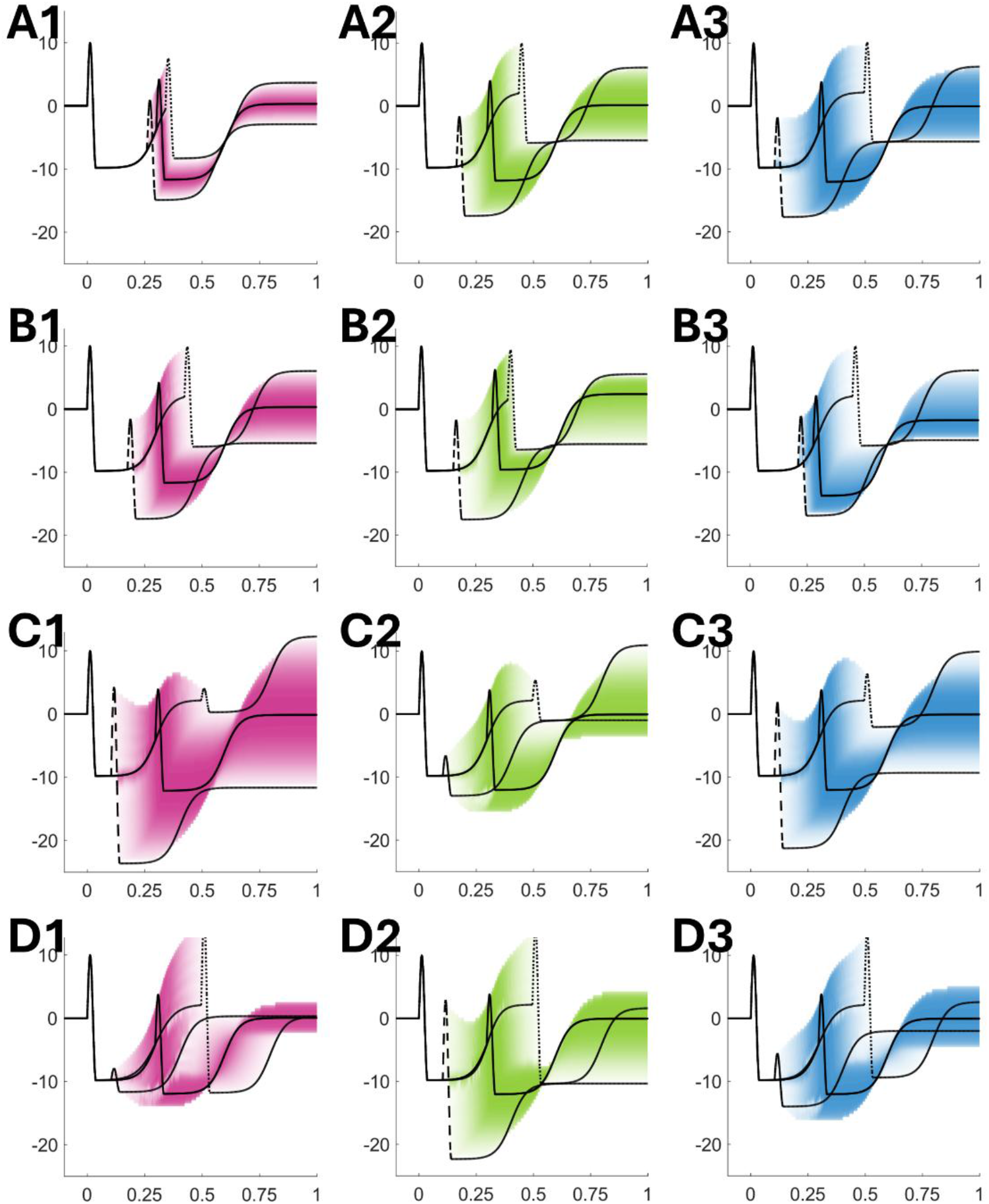
Simulation classes: Rows correspond to four simulation classes (A–D), columns to simulation levels (1–3) within each class. In every panel, solid black trace represents ground-truth response generated as median of sampled parameters. Dashed line represents 5th percentile, dotted line 95th percentile response. Coloured shading depicts density of sampled responses. A—Gaussian onset jitter: Gaussian onset jitter applied to second event with standard deviations 25 ms (A1), 75 ms (A2) and 125 ms (A3). B—Skewed onset distributions: Event onsets drawn from Pearson Type VI distribution with skewness −1 (B1), 0 (B2) and +1 (B3). C—Amplitude-onset coupling: Power-law coupling between second burst amplitude and onset latency, with exponents 1 (C1), 2 (C2) and 3 (C3). D—Combined coupling: Combined coupling where both second burst amplitude and first recovery slope vary as power-law functions of second event onset latency, tested at exponents 1 (D1), 2 (D2) and 3 (D3).

#### B. Skewed onset variability (Figure 3B)

To capture asymmetric latencies observed in reaction-time experiments, we sampled event onsets from Pearson Type VI distributions (scaled F-distribution) with fixed standard deviation 100 ms, kurtosis 4.5, and skewness −1, 0 and +1, generating left-skewed, symmetric and right-skewed timing distributions.

#### C. Amplitude-onset coupling (Figure 3C)

Real neural responses often exhibit relationships between response strength and timing. We coupled second transient amplitude to the onset latency via power-law relationships (exponents 1, 2 or 3: linear, quadratic or cubic scaling). Underlying onset jitter was drawn from Gaussian distribution with 100 ms standard deviation.

#### D. Combined slope-onset and amplitude-onset coupling (Figure 3D)

We retained power-law coupling between second event amplitude and onset latency whilst introducing a second power-law dependency where the first transient recovery slope varied with second event onset. Onset jitter was Gaussian (σ = 100 ms). We tested linear, quadratic and cubic exponents for both coupling functions, examining how simultaneous timing-magnitude interactions impact alignment performance.

Each simulation type was repeated 250 times with 60 trials each. Inter-trial variability was imposed as described, and temporally correlated 1/*f* noise added to achieve SNR = 0.3 (defined as signal RMS divided by noise RMS).

We quantified alignment accuracy using standardised root-mean-square error (sRMSE) between the ERW-recovered average and noise-free ground truth. Ground truth was defined by generating the response from the template at median event onset, yielding an idealised waveform without jitter or noise. sRMSE measures pointwise deviation of the estimated average from ground truth, scaled by ground truth standard deviation. We computed sRMSE over 200–800 ms post-stimulus, capturing >95% of event-related signal variance and concentrating error measurement on periods most sensitive to temporal misalignment.

All simulations used DST order K = 8 to adequately constrain warping trajectories and ensure smooth deformations. Progressive alignment employed 18 intermediate templates, providing sufficient granularity to avoid local minima. Optimisation tolerances (step and objective function) of 5 × 10^−6^ balanced convergence precision against computational efficiency.

### 2.3 Empirical Validation

Whilst simulation studies demonstrate ERW can recover known ground-truth signals under controlled conditions, empirical validation with real neurophysiological data is essential to establish construct validity and practical utility. We address three objectives. First, we assess whether ERW-aligned event-related potentials successfully recover sequential responses in multichannel data, focusing on second-event recovery accuracy as a stringent test of alignment efficacy. Second, we verify that temporal realignment preserves intrinsic lead-lag relationships between channels, ensuring suitability for subsequent causal and connectivity inference. Third, we examine whether distance-weighted averaging, down-weighting trials requiring substantial temporal correction, further improves signal recovery. Together, these analyses establish whether ERW is appropriate for neuroimaging applications characterised by experimental jitter, correlated noise and multichannel recordings.

#### 2.3.1 Dataset

We analysed an open-access auditory go/no-go dataset (https://openneuro.org/datasets/ds003690) comprising 64-channel EEG from 75 participants (36 younger adults: 23 ± 3 years, 29 female; 39 older adults: 60 ± 5 years, 31 female). Participants had normal or corrected vision and hearing with no neurological or psychiatric history. The task involved cued auditory choice trials with variable cue-to-target intervals, making it well-suited for evaluating ERW under naturalistic timing variability.

Each trial presented an anticipatory cue followed after a variable delay by either a go signal (250 ms, 1700 Hz tone, 80 trials/run) or a no-go signal (250 ms, 1300 Hz tone, 20 trials/run) at approximately 67 dB(A). Participants pressed a key for go trials and withheld responses for no-go trials. Commission errors and go responses exceeding 700 ms triggered 250 ms, 1000 Hz feedback tones 1200 ms post-stimulus. Critically, cue-to-target intervals were randomised using truncated exponential distributions spanning 1.5–4.1 s, whilst post-stimulus intervals ranged 5.2–15.5 s, yielding total inter-trial intervals of 6.7–19.6 s. This large temporal variability provides naturalistic testing of ERW’s ability to recover event-related responses when sequential event timing varies substantially across trials.

EEG was recorded at 500 Hz using a 64-channel International 10-20 system (reference CPz, ground FPz, forehead electrodes excluded). Each participant completed two approximately 8-minute runs with self-paced breaks. We analysed correctly performed go and no-go trials only, discarding behavioural errors and artefact-contaminated epochs.

#### 2.3.2 Preprocessing

Continuous EEG was imported into SPM25 (Wellcome Centre for Human Neuroimaging, London) and band-pass filtered (4th-order Butterworth, 0.1–30 Hz) to attenuate drift and high-frequency noise whilst preserving ERP morphology. Data were downsampled to 200 Hz, balancing fidelity against computational efficiency. We epoched recordings from −1000 to +6000 ms relative to trial onset (anticipatory cue), encompassing both cue and target events, without baseline correction to avoid distorting temporal alignment.

We excluded epochs containing behavioural errors (no-go commissions, go omissions, responses >700 ms) and rejected epochs that exhibited |z| > 7 relative to within-channel distribution, removing movement and muscle artefacts. Channels were then re-referenced to the common average.

To assess weighted averaging in the absence of ground truth, we defined “go (select)” by restricting go trials to cue-to-target intervals within ±100 ms of the sample median (1680 ms). Reduced temporal variability should enhance target-locked ERP fidelity, providing an approximate benchmark. However, this is not the gold standard: temporal restriction reduces trial counts, introduces selection bias and does not eliminate residual variability. Consequently, convergence between distance-weighted ERW averages and go (select) provides suggestive rather than definitive evidence.

#### 2.3.3 Event-Related Warping Procedure

We implemented temporal alignment using the ERW toolbox for SPM25 (Appendix lists complete functions). The procedure comprised four stages: design specification, template construction, warp trajectory estimation and signal alignment.

##### Design specification (spm_eeg_erw_design)

For each participant, we constructed design matrices encoding two conditions: “go (trial)” and “no-go (trial)”. Each comprising two sequential events: trial-onset (anticipatory cue) and target (go or no-go imperative signal). Event onsets were set to recorded stimulus times. Each event was represented as Gaussian “bump” function with dispersion σ = 50 ms, providing smooth continuous representation of event timing. Temporal warping functions were parameterised using 8th-order DST basis with Tukey tapering (taper ratio 0.1), minimising warping at boundaries where minimal temporal jitter was expected.

##### Template construction (spm_eeg_erw_template)

For each participant, we computed subject-specific template target end-points by averaging event onset times across conditions (go and no-go), yielding within-subject consensus timing. We then derived group-level templates target end-points by averaging subject-specific event times across all participants, establishing a common temporal reference reflecting the central tendency of event timing across the group. For each participant, we generated 18 intermediate templates linearly interpolating from the original (participant-specific, condition-specific) event times to the final group-level target template.

##### Warp trajectory estimation (spm_eeg_erw_estimate)

We estimated subject-specific and condition-specific warp trajectories by minimising mean squared error between successive templates in the interpolation sequence. For each trial, the template at step *s* was lifted to continuous time using parabolic interpolation, and a smooth monotonic warping function τ(t) (represented via DST basis) was optimised to align this template to the template at step *s* + 1. Optimisation employed the BFGS quasi-Newton algorithm using analytic gradients derived in Section 1.1. We set the step tolerance to 5 × 10^−6^ and the objective function tolerance to 5 × 10^−6^. This process was repeated across all 18 interpolation steps, progressively refining warp trajectories until each trial’s event structure aligned to the group template.

##### Signal alignment (spm_eeg_erw_warp)

Epoched EEG data were temporally aligned by applying estimated warp trajectories τ to each trial. Trials were resampled on deformed timelines using modified Akima interpolation.

##### Trial averaging (spm_eeg_erw_average)

We computed trial-averaged ERPs using conventional (unweighted) averaging and distance-weighted averaging. For distance-weighted averaging, each trial received a weight based on the total temporal deviation from the target template.

#### 2.3.4 Validation

We generated reference signals by computing event-related responses for target events (go or no-go) using conventional epoching and averaging. Alongside go (select) responses to appraise weighting procedures. We tested four weights for trial averaging: *κ* ∈ {0, 1, 2, 3}, where \k = 0 collapses to uniform (unweighted) averaging.

##### Event-related potential similarity

For each participant, condition (go vs no-go) and weighting parameter κ, we calculated a standardised root-mean-square error (sRMSE) between the warped ERP and reference ERP over 0–500 ms post-target using only channels marked good. Specifically, for each channel:

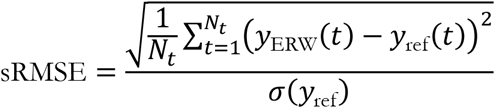

This standardisation accounts for signal amplitude differences across channels and participants. We averaged the sRMSE across all good channels, obtaining single summary values per participant, condition and weighting scheme. A lower sRMSE indicates greater similarity between ERW-aligned average and reference, interpreted as evidence that ERW successfully reduced trial-to-trial variability without distorting underlying ERP morphology. We compared sRMSE values across weighting schemes and between go and no-go conditions.

##### Inter-sensor timing preservation

To verify that temporal realignment did not alter intrinsic lead-lag relationships among channels, we conducted cross-covariance analysis comparing ERW-aligned and reference data. For every unique pair of good EEG channels, we computed normalised cross-correlation functions over ±200 ms lags within 0–500 ms post-target window For each channel pair, we identified the lag where absolute cross-correlation reached maximum, denoted *L*_ref_ for reference data and *L*_ERW_ for ERW-aligned data.

To assess whether ERW preserved temporal relationships, we quantified lag shift magnitude as normalised root-mean-square lag error:

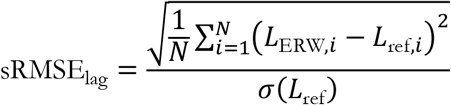

where sum runs over all channel pairs and *σ*(*L*_ref_) is the standard deviation of reference lags across pairs. We expected ERW should minimally perturb inter-channel lag structure, manifesting as low normalised lag errors (<0.1). Systematic deviations would suggest ERW introduces artefactual timing shifts, undermining physiological interpretability.

## 3 Results

### 3.1 Simulation

We validated ERW’s ability to recover known temporal structure using synthetic data incorporating realistic neurophysiological variability. Four simulation classes systematically manipulated trial-to-trial timing relationships: (A) Gaussian onset jitter, (B) skewed latency distributions, (C) amplitude-latency coupling, and (D) Amplitude-latency and slope-latency coupling. Each class comprised 250 iterations of 60 trials with temporally correlated 1/f noise (SNR = 0.3). We quantified recovery accuracy using standardised root-mean-square error (sRMSE) between ERW-recovered averages and noise-free ground truth, computed over 200– 800 ms post-stimulus. The reference provides a baseline representing the sMRSE of a non-jittered trial averaged canonical response with additive noise (sRMSE = 0.296). Results are compared between conventional averaging (CA) and distance-weighted averaging with sharpness parameters *κ* ∈ 1,2,3.

#### 3.1.1 Simulation A: Gaussian Onset Variability

Gaussian jitter in second-event timing at *σ* ∈ 25,75,125*ms* tested ERW’s performance under symmetric, increasing temporal dispersion (**Figure 4, Table 1**). At low variability (25 ms), CA achieved near-reference performance (0.306, 95%CI [0.297, 0.316]), with weighting offering no benefit and *κ* = 1 slightly deleterious (−6.4%). At moderate variability (75 ms), CA yielded 0.344, 95%CI [0.332, 0.356], and optimal weighting (*κ* = 2) reduced error by 1.6%, 95%CI [0.8%, 2.5%]. Time-resolved traces showed residual error slightly above the noise floor around the second event (**Figure 4 B2**). High variability (125 ms) revealed clearer benefits for weighting: CA produced 0.376, 95%CI [0.361, 0.393], whilst *κ* = 1 achieved 3.5%, 95%CI [1.4%, 5.6%] improvement. Enhanced gains appeared in the late post-stimulus window (600–800 ms, **Figure 4 C2**), where weighted averaging effectively down-weighted trials requiring large temporal corrections that introduced secondary distortions. Representative traces (**Figure 4, row 3**) demonstrated close correspondence with ground truth across all levels when optimal parameters were employed. Overall, ERW aligned well across skew regimes, and weighting improved recovery consistently, with kappa 1 preferred.

**Table 1:** Gaussian onset variability: sRMSE for conventional averaging (CA) and distance-weighted averaging (WA) at *κ* ∈ {1,2,3}, with second-event onset jitter *σ* ∈ {25,75,125} ms. Values represent means and 95% confidence intervals across 250 simulations (60 trials each; SNR = 0.3). Pairwise improvement quantifies within-simulation percentage sRMSE reduction relative to CA; asterisks denote optimal weighting scheme for each jitter level.

|  | Averaging Method | sRMSE Mean [95% CI] | SD | Pairwise improvement in sRMSE compared with CA [95% CI] |
| --- | --- | --- | --- | --- |
| $\sigma: 25$ ms | CA | 0.306 [0.297, 0.316] | 0.048 | — |
| | WA($\kappa:1$) | 0.324 [0.315, 0.334] | 0.048 | -6.38% [-8.15%, -4.63%] Higher |
| | WA($\kappa:2$) | 0.307 [0.298, 0.316] | 0.047 | -0.42% [-1.05%, 0.21%] Higher |
| | WA($\kappa:3$) | 0.306 [0.297, 0.315] | 0.047 | 0.08% [-0.23%, 0.39%] Lower* |
| $\sigma: 75$ ms | CA | 0.344 [0.332, 0.356] | 0.060 | — |
| | WA($\kappa:1$) | 0.343 [0.332, 0.354] | 0.055 | -0.67% [-3.00%, 1.77%] Higher |
| | WA($\kappa:2$) | 0.337 [0.326, 0.348] | 0.056 | 1.63% [0.82%, 2.49%] Lower* |
| | WA($\kappa:3$) | 0.340 [0.329, 0.351] | 0.058 | 1.00% [0.61%, 1.42%] Lower |
| $\sigma: 125$ ms | CA | 0.376 [0.361, 0.393] | 0.083 | — |
| | WA($\kappa:1$) | 0.360 [0.346, 0.375] | 0.075 | 3.51% [1.35%, 5.59%] Lower* |
| | WA($\kappa:2$) | 0.365 [0.350, 0.381] | 0.079 | 2.89% [2.20%, 3.59%] Lower |
| | WA($\kappa:3$) | 0.370 [0.355, 0.387] | 0.081 | 1.54% [1.22%, 1.87%] Lower |

**Figure 4:**
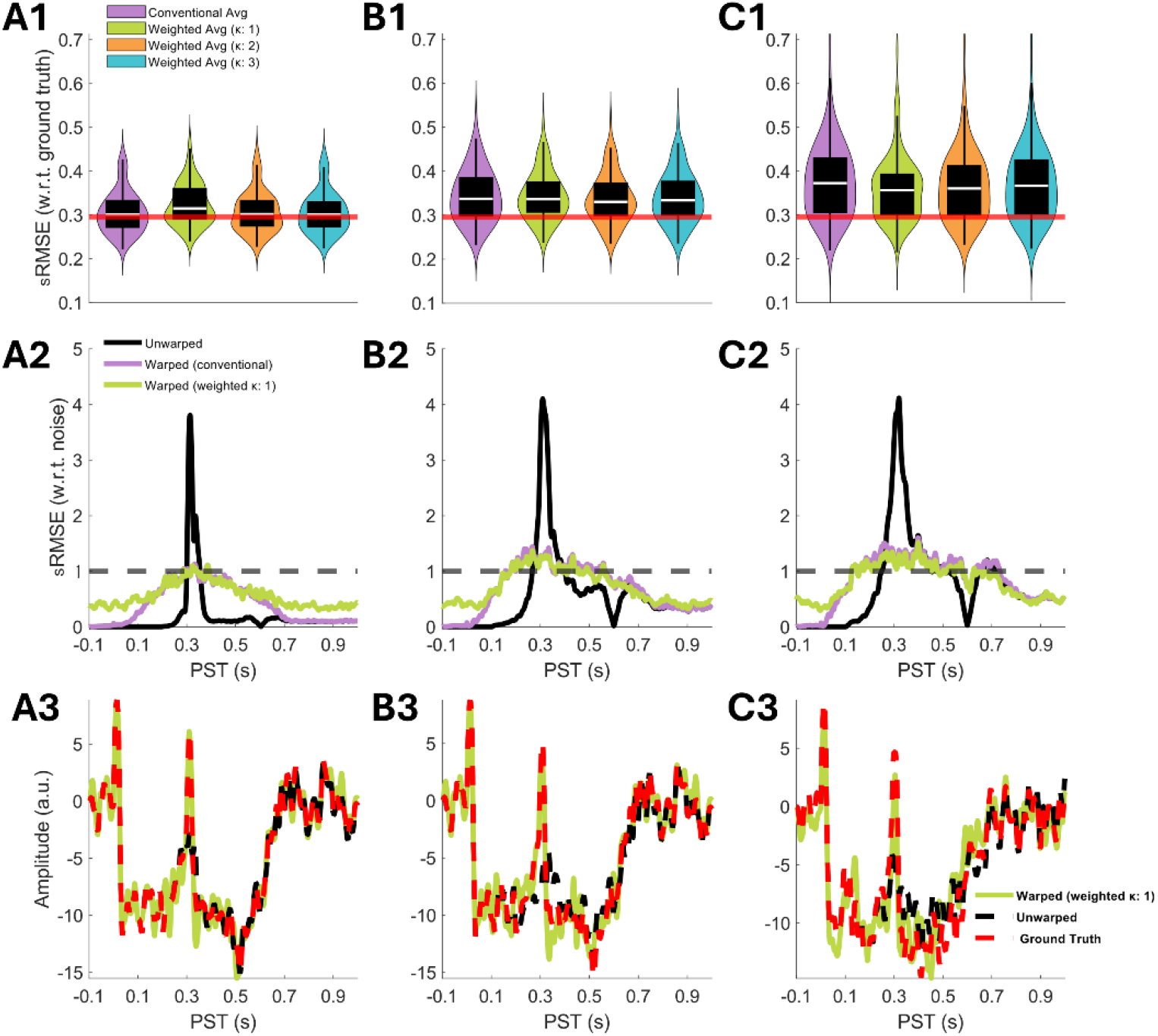
Gaussian onset variability outcomes: Columns indicate second-event jitter dispersion: A, σ = 25 ms; B, σ = 75 ms; C, σ = 125 ms. Row 1: sRMSE distributions normalised by ground-truth standard deviation (200–800 ms window). Violin plots show median (white line), interquartile range (thick bar), and extreme non-outlier values (whiskers). Red horizontal line marks non-jittered reference (0.296). Row 2: Time-resolved sRMSE normalised by noise standard deviation and averaged across simulations; dashed line at 1.0 marks noise floor. Row 3: Representative recovered responses from median-performing iteration, comparing warped weighted average (green), unwarped response (black), and ground truth (red dashed).

#### 3.1.2 Simulation B: Skewed Latency Distributions

To model reaction-time-like asymmetries, second-event onsets were drawn from Pearson Type VI distributions (*σ* = 100 ms, kurtosis = 4.5) with skew ∈ {−1, 0, +1} (**Figure 5, Table 2**). Symmetric timing (skew = 0) resembled moderate Gaussian jitter: CA yielded 0.307, 95%CI [0.295, 0.319], and *κ* = 1 reduced error by 5.0%, 95%CI [3.6%, 6.4%]. Time-resolved error remained at or below the noise floor (**Figure 5 A2**), demonstrating effective recovery despite substantial temporal variability. Left-skewed timing (skew = −1, early-biased outliers) produced comparatively elevated localisation error; however, ERW still offered a high level of signal recovery: CA produced 0.380, 95%CI [0.365, 0.395], with a 10.1% 95%CI [8.6%, 11.6%] improvement via *κ* = 1. The elevated error localised to the initial burst (200–350 ms, **Figure 5 B2**). Distance weighting yielded substantial late-window improvements (400–800 ms) below the noise floor. Right-skewed timing (skew = +1, late-biased outliers) facilitated superior recovery: CA achieved 0.302, 95%CI [0.293, 0.311], with *κ* = 1 further reducing error by 10.1%, 95%CI [8.7%, 11.5%] to 0.271, 95%CI [0.262, 0.280] approaching reference levels. The weighted average remained close to or below the noise floor for most of the analysis window (**Figure 5 C2**). Overall, ERW aligned well across skew regimes, and weighting improved recovery consistently, with *κ*=1 preferred.

**Table 2:** Skewed latency distributions: sRMSE for conventional averaging (CA) and distance-weighted averaging (WA) at *κ* ∈ {1,2,3}, with second-event onsets drawn from Pearson Type VI distributions (*σ* = 100 ms, kurtosis = 4.5) at skew ∈ {−1,0, +1}. Values represent means and 95% confidence intervals across 250 simulations (60 trials each; SNR = 0.3). Pairwise improvement quantifies within-simulation percentage sRMSE reduction relative to CA; asterisks denote optimal weighting scheme for each skewness level.

|  | Averaging Method | sRMSE Mean [95% CI] | SD | Pairwise improvement in sRMSE compared with CA [95% CI] |
| --- | --- | --- | --- | --- |
| **Skew(0)** | CA | 0.307 [0.295, 0.319] | 0.061 | — |
| | WA($\kappa$:1) | 0.292 [0.280, 0.304] | 0.061 | 4.96% [3.55%, 6.36%] Lower* |
| | WA($\kappa$:2) | 0.296 [0.284, 0.308] | 0.061 | 3.68% [3.12%, 4.25%] Lower |
| | WA($\kappa$:3) | 0.301 [0.289, 0.313] | 0.061 | 1.99% [1.71%, 2.28%] Lower |
| **Skew(-1)** | CA | 0.380 [0.365, 0.395] | 0.078 | — |
| | WA($\kappa$:1) | 0.339 [0.327, 0.352] | 0.064 | 10.08% [8.58%, 11.59%] Lower* |
| | WA($\kappa$:2) | 0.359 [0.346, 0.374] | 0.071 | 5.25% [4.65%, 5.84%] Lower |
| | WA($\kappa$:3) | 0.369 [0.355, 0.384] | 0.075 | 2.67% [2.38%, 2.97%] Lower |
| **Skew(1)** | CA | 0.302 [0.293, 0.310] | 0.046 | — |
| | WA($\kappa$:1) | 0.271 [0.262, 0.280] | 0.044 | 10.09% [8.70%, 11.45%] Lower* |
| | WA($\kappa$:2) | 0.284 [0.275, 0.293] | 0.045 | 6.01% [5.38%, 6.63%] Lower |
| | WA($\kappa$:3) | 0.292 [0.283, 0.301] | 0.045 | 3.22% [2.89%, 3.55%] Lower |

**Figure 5:**
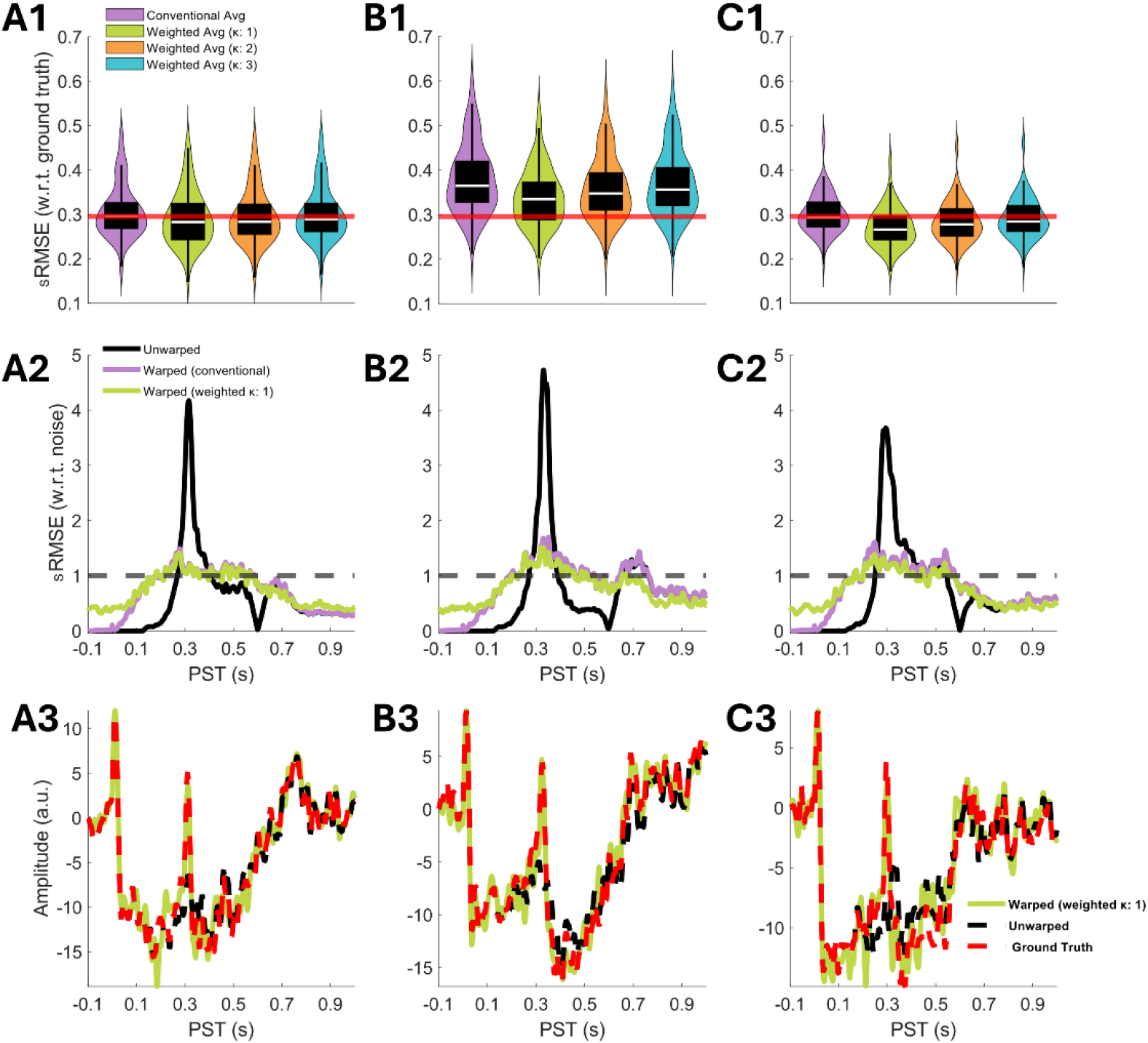
Skewed latency distribution outcomes: Columns indicate second-event onset skewness: A, skew = 0; B, skew = −1; C, skew = +1. Row 1: sRMSE distributions normalised by ground-truth standard deviation (200–800 ms window). Violin plots show median (white line), interquartile range (thick bar), and extreme non-outlier values (whiskers). Red horizontal line marks non-jittered reference (0.296). Row 2: Time-resolved sRMSE normalised by noise standard deviation and averaged across simulations; dashed line at 1.0 marks noise floor. Row 3: Representative recovered responses from median-performing iteration, comparing warped weighted average (green), unwarped response (black), and ground truth (red dashed).

#### 3.1.3 Simulation C: Amplitude-Latency Coupling

Power-law coupling between second-event amplitude and onset latency (exponents 1, 2, 3; Gaussian onset jitter *σ* = 100 ms) mimicked time-dependent amplitude responses (**Figure 6, Table 3**). Linear coupling produced CA sRMSE of 0.328, 95%CI [0.319, 0.336], with *κ* = 1 yielding 7.0%, 95%CI [5.6%, 8.3%] improvement. Quadratic coupling showed strong benefits for weighted averaging: CA achieved 0.342, 95%CI [0.330, 0.355], and *κ* = 1 reduced error by 13.2%, 95%CI [11.6%, 14.7%], the largest improvement observed across all simulations. Time-resolved analysis revealed sustained late-window improvements (500–800 ms, **Figure 6 B2**), suggesting weighted averaging effectively compensated for amplitude-timing co-variation that temporal alignment alone can not correct for. Cubic coupling also demonstrated a clear benefit for weighted averaging (*κ* = 1) (7.3%, 95%CI [5.9%, 8.6%]). Overall, ERA recovered responses close to the ground truth, and weighting delivered additional gains across profiles, strongest for quadratic coupling, with *κ* = 1 preferred. Time-resolved nRMSE remained at or marginally above the noise floor across all coupling strengths, with the largest reductions concentrated around the second event (300–500 ms). Representative traces (**Figure 6, row 3**) demonstrated tight correspondence with ground truth.

**Table 3:** Amplitude-latency coupling: sRMSE for conventional averaging (CA) and distance-weighted averaging (WA) at *κ* ∈ {1,2,3}, with second-event amplitude coupled to onset latency via power-law relationships (exponents 1, 2, 3; Gaussian onset jitter *σ* = 100 ms). Values represent means and 95% confidence intervals across 250 simulations (60 trials each; SNR = 0.3). Pairwise improvement quantifies within-simulation percentage sRMSE reduction relative to CA; asterisks denote optimal weighting scheme for each coupling strength.

|  | Averaging Method | sRMSE |  | Pairwise improvement in sRMSE compared with CA [95% CI] |
| --- | --- | --- | --- | --- |
|  |  | Mean [95% CI] | SD |  |
| linear | CA | 0.328 [0.319, 0.336] | 0.045 | — |
| | WA($\kappa$:1) | 0.304 [0.296, 0.312] | 0.041 | 6.98% [5.58%, 8.33%] Lower* |
| | WA($\kappa$:2) | 0.312 [0.303, 0.320] | 0.042 | 4.82% [4.25%, 5.40%] Lower |
| | WA($\kappa$:3) | 0.319 [0.310, 0.327] | 0.044 | 2.62% [2.32%, 2.93%] Lower |
| quadratic | CA | 0.342 [0.329, 0.355] | 0.065 | — |
| | WA($\kappa$:1) | 0.294 [0.285, 0.303] | 0.046 | 13.18% [11.58%, 14.74%] Lower* |
| | WA($\kappa$:2) | 0.319 [0.307, 0.330] | 0.058 | 6.73% [6.15%, 7.30%] Lower |
| | WA($\kappa$:3) | 0.330 [0.318, 0.343] | 0.062 | 3.42% [3.13%, 3.71%] Lower |
| cubic | CA | 0.327 [0.316, 0.338] | 0.055 | — |
| | WA($\kappa$:1) | 0.302 [0.293, 0.311] | 0.048 | 7.29% [5.93%, 8.64%] Lower* |
| | WA($\kappa$:2) | 0.313 [0.303, 0.323] | 0.052 | 4.30% [3.84%, 4.77%] Lower |
| | WA($\kappa$:3) | 0.320 [0.309, 0.330] | 0.053 | 2.25% [2.03%, 2.48%] Lower |

**Figure 6:**
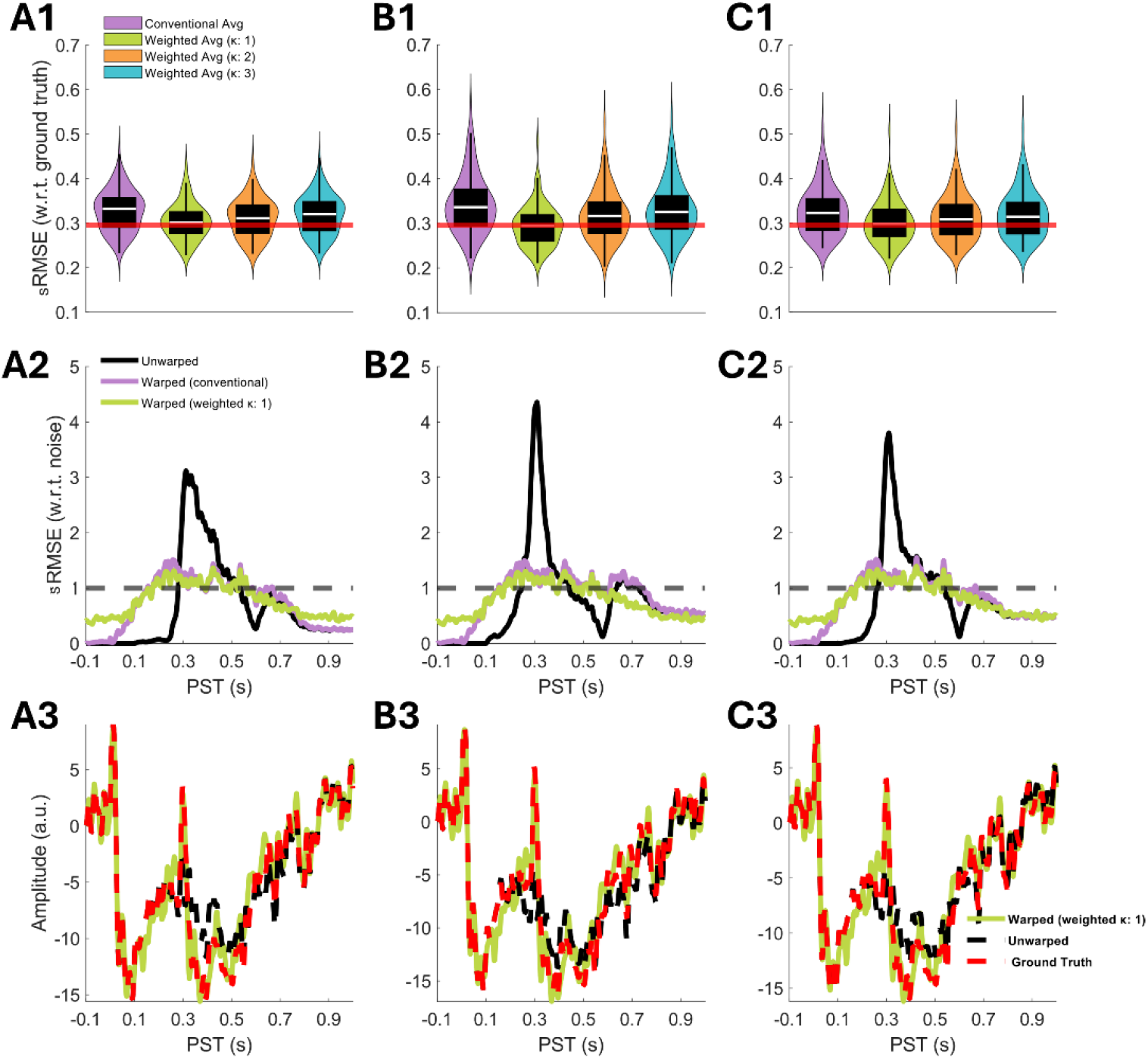
Amplitude-latency coupling outcomes. Columns indicate power-law coupling strength between second-event amplitude and onset: A, linear; B, quadratic; C, cubic. Row 1: sRMSE distributions normalised by ground-truth standard deviation (200–800 ms window). Violin plots show median (white line), interquartile range (thick bar), and extreme non-outlier values (whiskers). Red horizontal line marks non-jittered reference (0.296). Row 2: Time-resolved sRMSE normalised by noise standard deviation and averaged across simulations; dashed line at 1.0 marks noise floor. Row 3: Representative recovered responses from median-performing iteration, comparing warped weighted average (green), unwarped response (black), and ground truth (red dashed).

#### 3.1.4 Simulation D: Amplitude-Latency and Slope-Latency Coupling

Combined power-law dependencies of second-event amplitude and first-event recovery slope, both coupled to second-event onset, tested ERW under complex, multivariate timing-morphology relationships (exponents 1, 2, 3; Gaussian onset jitter *σ* = 100ms) (**Figure 7, Table 4**). Linear coupling showed 5.8%, [4.9%, 6.8%] improvement with *κ* = 1; quadratic coupling achieved 12.7%, 95%CI [11.7%, 13.7%] reduction; cubic coupling yielded 6.7%, 95%CI [5.8%, 7.5%] benefit. Time-resolved traces showed aligned responses near the noise floor throughout the analysis window, with sustained late-period benefits most prominent in quadratic coupling (**Figure 7 B2**). Representative waveforms (**Figure 7, row 3**) closely matched ground truth across all coupling regimes, demonstrating successful disentanglement of timing and morphology despite their interdependence. ERW with weighted averaging robustly handles interdependent temporal-morphological variation without substantial performance degradation, validating its applicability to real neural data where multiple response properties co-vary with timing.

**Table 4:** Multi-parameter coupling: sRMSE for conventional averaging (CA) and distance-weighted averaging (WA) at *κ* ∈ {1,2,3}, with second-event amplitude and first-event recovery slope both coupled to second-event onset via power-law relationships (exponents 1, 2, 3; Gaussian onset jitter *σ* = 100 ms). Values represent means and 95% confidence intervals across 250 simulations (60 trials each; SNR = 0.3). Pairwise improvement quantifies within-simulation percentage sRMSE reduction relative to CA; asterisks denote optimal weighting scheme for each coupling strength.

|  | Averaging Method | sRMSE Mean [95% CI] | SD | Pairwise improvement in sRMSE compared with CA [95% CI] |
| --- | --- | --- | --- | --- |
| linear | CA | 0.329 [0.322, 0.335] | 0.051 | — |
| | WA($\kappa$:1) | 0.308 [0.302, 0.314] | 0.046 | 5.83% [4.85%, 6.79%] Lower* |
| | WA($\kappa$:2) | 0.315 [0.309, 0.321] | 0.048 | 4.17% [3.77%, 4.57%] Lower |
| | WA($\kappa$:3) | 0.321 [0.315, 0.327] | 0.049 | 2.29% [2.09%, 2.49%] Lower |
| quadratic | CA | 0.346 [0.337, 0.356] | 0.076 | — |
| | WA($\kappa$:1) | 0.298 [0.292, 0.305] | 0.053 | 12.69% [11.65%, 13.71%] Lower* |
| | WA($\kappa$:2) | 0.323 [0.314, 0.331] | 0.068 | 6.53% [6.13%, 6.92%] Lower |
| | WA($\kappa$:3) | 0.334 [0.326, 0.344] | 0.072 | 3.32% [3.13%, 3.52%] Lower |
| cubic | CA | 0.328 [0.320, 0.337] | 0.067 | — |
| | WA($\kappa$:1) | 0.305 [0.298, 0.312] | 0.057 | 6.65% [5.77%, 7.51%] Lower* |
| | WA($\kappa$:2) | 0.315 [0.307, 0.323] | 0.064 | 4.02% [3.70%, 4.35%] Lower |
| | WA($\kappa$:3) | 0.321 [0.314, 0.330] | 0.065 | 2.12% [1.96%, 2.28%] Lower |

**Figure 7:**
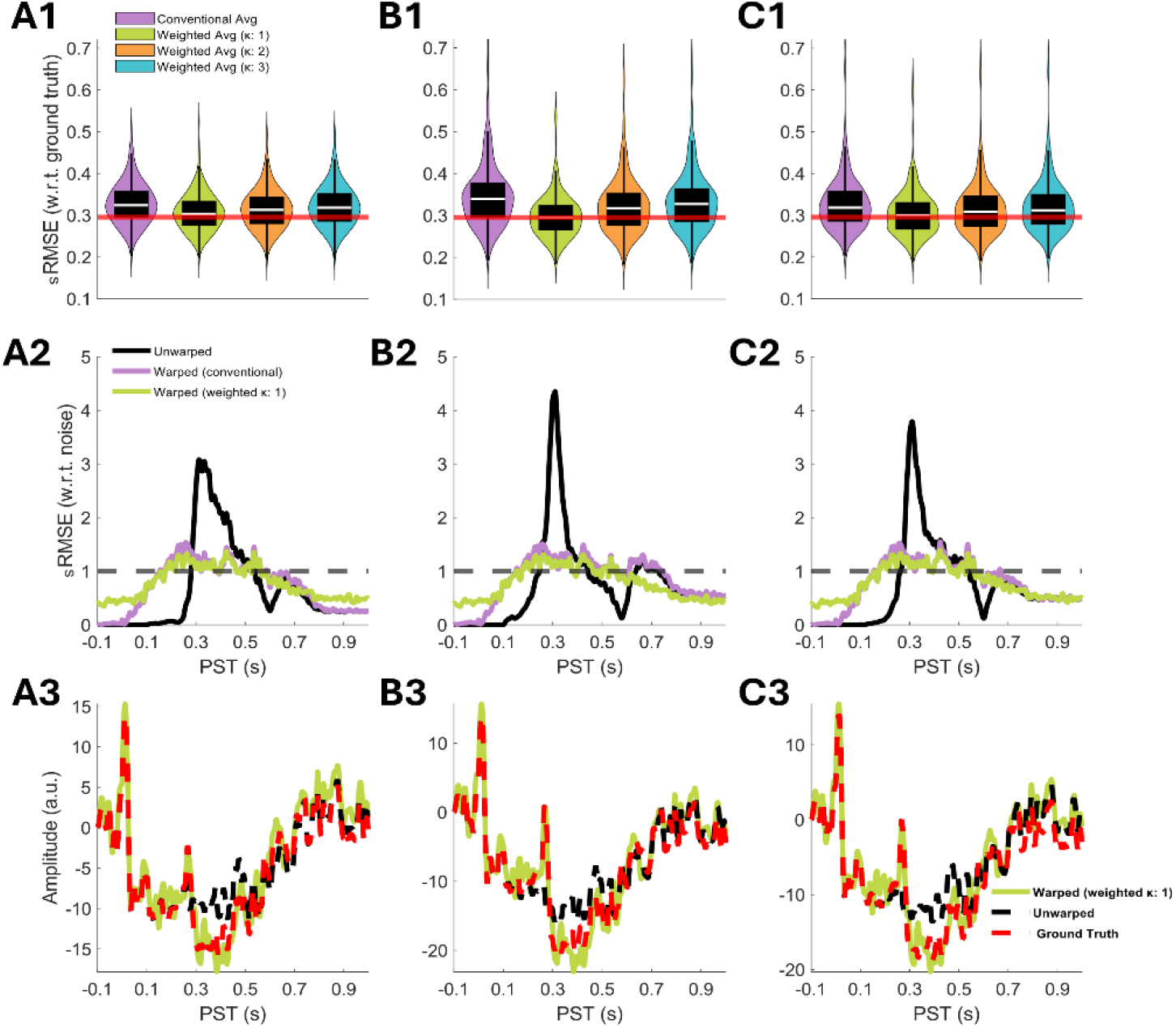
Multi-parameter coupling outcomes. Columns indicate power-law coupling strength, with both second-event amplitude and first-event recovery slope coupled to second-event onset: A, linear; B, quadratic; C, cubic. Row 1: sRMSE distributions normalised by ground-truth standard deviation (200–800 ms window). Violin plots show median (white line), interquartile range (thick bar), and extreme non-outlier values (whiskers). Red horizontal line marks non-jittered reference (0.296). Row 2: Time-resolved sRMSE normalised by noise standard deviation and averaged across simulations; dashed line at 1.0 marks noise floor. Row 3: Representative recovered responses from median-performing iteration, comparing warped weighted average (green), unwarped response (black), and ground truth (red dashed).

#### 3.1.5 Summary

ERW demonstrated robust recovery of temporally jittered responses across diverse neurophysiological variability regimes, with all aligned averages substantially outperforming unaligned baselines regardless of jitter characteristics. Time-resolved error concentrated around manipulated events (200–500 ms) whilst preserving pre-stimulus baselines, confirming genuine temporal realignment. Distance-weighted averaging provided minimal benefit at low variability (*σ* ≤ 25ms) but yielded systematic improvements of 5–13% under realistic experimental jitter levels (*σ* ≥ 100ms), with maximal benefits (≈13% error reduction) observed under quadratic amplitude-latency coupling, arguably the most ecologically valid scenario, suggesting weighted averaging is particularly valuable when response intensity and timing co-vary systematically. The sharpness parameter *κ* = 1 proved optimal across most manipulations, balancing noise suppression against over-aggressive trial exclusion.

### 3.2 Empirical

We validated ERW’s performance on real neurophysiological data using an open-access auditory go/no-go EEG dataset. The task incorporated variable cue-to-target intervals (1.5 to 4.1 s), providing large temporal variability for evaluating ERW under experimental conditions. We assessed whether ERW-aligned event-related potentials recovered sequential response by preserving cue-onset responses and target-onset responses, and whether temporal realignment preserved intrinsic interchannel lead-lag relationships.

We quantified ERP recovery using sRMSE between ERW-aligned averages and epoched reference signals computed over 0 to 500 ms post-target onset. Intechannel timing preservation was assessed via cross-covariance analysis comparing lag structures between ERW-aligned and reference data. Results are reported separately for go trials, no-go trials, and temporally restricted go trials where cue-to-target intervals fell within ± 100 ms of the sample median (go select). Distance-weighted averaging with sharpness parameters *κ* ∈ 1,2,3 was examined; however, the absence of ground truth in empirical data limits interpretation of optimal weighting to illustrative comparisons against the epoched reference.

#### 3.2.1 Event-Related Responses

ERW-aligned averages demonstrated successful recovery of target-locked responses relative to conventionally epoched references (**Figure 8, Table 5**).

**Table 5:** Empirical ERP data: sRMSE for conventional averaging (CA) and distance-weighted averaging (WA) at *κ* ∈ {1,2,3}, across three ERP datasets. Values represent means and 95% confidence intervals. Pairwise improvement quantifies within-subject percentage sRMSE reduction relative to CA; asterisks denote optimal weighting scheme for each dataset.

|  |  |  |  |  |
| --- | --- | --- | --- | --- |
| **Go** | CA | 0.239 [0.219, 0.259] | 0.062 | — |
| | WA($\kappa:1$) | 0.389 [0.351, 0.429] | 0.123 | -64.00% [-73.92%, -54.45%] Higher |
| | WA($\kappa:2$) | 0.286 [0.261, 0.312] | 0.078 | -20.89% [-27.13%, -15.08%] Higher |
| | WA($\kappa:3$) | 0.255 [0.234, 0.275] | 0.066 | -7.28% [-10.54%, -4.26%] Higher |
| **No-Go** | CA | 0.333 [0.288, 0.381] | 0.144 | — |
| | WA($\kappa:1$) | 0.512 [0.443, 0.588] | 0.224 | -58.91% [-72.27%, -46.93%] Higher |
| | WA($\kappa:2$) | 0.375 [0.324, 0.428] | 0.162 | -13.83% [-18.32%, -9.52%] Higher |
| | WA($\kappa:3$) | 0.344 [0.298, 0.396] | 0.151 | -3.54% [-5.74%, -1.53%] Higher |
| **Go(Select)** | CA | 0.511 [0.460, 0.566] | 0.163 | — |
| | WA($\kappa:1$) | 0.364 [0.322, 0.410] | 0.135 | 28.94% [25.43%, 32.48%] Lower* |
| | WA($\kappa:2$) | 0.448 [0.403, 0.498] | 0.147 | 12.30% [10.49%, 14.08%] Lower |
| | WA($\kappa:3$) | 0.476 [0.429, 0.528] | 0.153 | 6.88% [5.84%, 7.93%] Lower |

**Figure 8:**
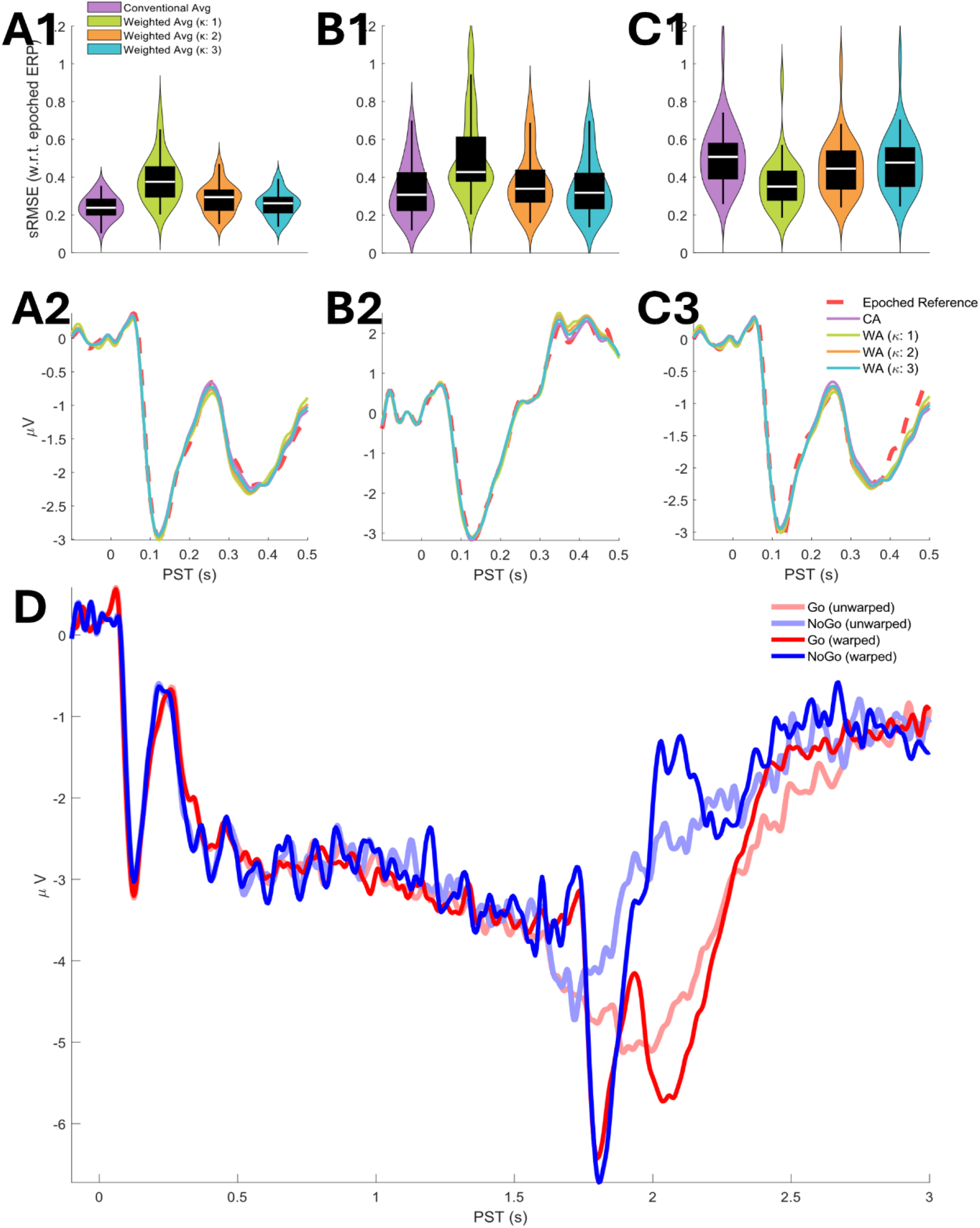
Event-related potential recovery and distance-weighted averaging validation. Columns indicate trial type: A, go trials B, no-go trials; C, go trials with restricted cue-to-target intervals. Row 1: sRMSE distributions comparing conventional averaging (purple) and weighted averaging at *κ* ∈ {1, 2, 3} (yellow-green, orange, cyan) against epoched reference (0 to 500 ms post-target). Violin plots show median (white line), interquartile range (thick bar), and extreme non-outlier values (whiskers). Lower nRMSE indicates greater similarity to reference. Row 2: Grand-average target-locked ERPs overlaying conventional and weighted averages with epoched reference (red dashed). D: Extended time-course validation (−100 to 3000 ms post-target). Unwarped go (light red) and no-go (light blue) exhibit temporal smearing; warped counterparts (dark red, dark blue) show sharpened target-locked components whilst preserving cue-related activity (0 to 100 ms).

For go and no-go trials, conventional averaging (CA) achieved sRMSE of 0.239, 95%CI [0.219, 0.259] and 0.333, 95%CI [0.288, 0.381], respectively. Distance-weighted averaging systematically increased error relative to CA across both conditions, with deviations ranging from 3.5% to 64.0% depending on the sharpness parameter. Grand-average waveforms (**Figure 8 A2, B2**) showed all averaging schemes closely approximated epoched reference morphology across the 0 to 500 ms post-target window, with minimal visual differentiation between conventional and weighted approaches.

For go (select) trials, CA produced sRMSE of 0.511, 95%CI [0.460, 0.566]. Distance-weighted averaging reduced error systematically: *κ* = 1 achieved 28.9%, 95%CI [25.4%, 32.5%] reduction; *κ* = 2 yielded 12.3%, 95%CI [10.5%, 14.1%] reduction; *κ* = 3 produced 6.9%, 95%CI [5.8%, 7.9%] reduction. Grand-average waveforms (**Figure 8 C3**) showed weighted averages more closely tracked epoched reference, particularly during mid-to-late negativity (200 to 400 ms).

Extended time-course analysis (**Figure 8D**) spanning the entire sequential epoch −100 to 3000 ms demonstrated preservation of cue-related activity and recovery of target-locked responses. Warped traces showed sharpened target components concentrated around 1700 to 3000 ms with preservation of cue-related activity (0 to 1700 ms post-target).

#### 3.2.2 Channel Cross-Covariance

Inter-channel lag structure preservation was assessed by comparing cross-covariance matrices between ERW-aligned and epoched reference signals across channel pairs exhibiting correlation r>0.60 (**Figure 9, Table 6**).

**Table 6:** Empirical cross-covariance data: sRMSE for conventional averaging (CA) and distance-weighted averaging (WA) at *κ* ∈ {1,2,3}, across three ERP datasets using cross-covariance-based trial selection (r>0.60). Values represent means and 95% confidence intervals. Pairwise improvement quantifies within-subject percentage nRMSE reduction relative to CA; asterisks denote optimal weighting scheme for each dataset.

|  | Method Averaging | Mean [95% CI] sRMSE | SD | compared with CA [95% CI] Pairwise improvement in sRMSE |
| --- | --- | --- | --- | --- |
| **Go**<br>$r > 0.60$ | CA | 0.631 [0.554, 0.720] | 0.257 | — |
| | WA($\kappa:1$) | 0.849 [0.719, 0.998] | 0.429 | -47.14% [-73.20%, -23.54%] |
| | WA($\kappa:2$) | 0.731 [0.646, 0.827] | 0.279 | -25.64% [-42.77%, -11.15%] |
| | WA($\kappa:3$) | 0.654 [0.587, 0.730] | 0.217 | -9.93% [-20.87%, -1.07%] |
| **No-Go**<br>$r > 0.60$ | CA | 0.705 [0.627, 0.783] | 0.239 | — |
| | WA($\kappa:1$) | 0.871 [0.752, 0.992] | 0.374 | -35.42% [-69.12%, -11.10%] |
| | WA($\kappa:2$) | 0.795 [0.697, 0.902] | 0.318 | -22.34% [-53.75%, -2.58%] |
| | WA($\kappa:3$) | 0.709 [0.629, 0.797] | 0.264 | -1.39% [-6.46%, 3.39%] |
| **Go (select)**<br>$r > 0.60$ | CA | 1.123 [0.939, 1.367] | 0.658 | — |
| | WA($\kappa:1$) | 0.829 [0.708, 0.969] | 0.409 | 19.23% [8.46%, 29.92%] Lower* |
| | WA($\kappa:2$) | 0.916 [0.823, 1.010] | 0.284 | 4.71% [-9.31%, 16.32%] Lower |
| | WA($\kappa:3$) | 0.958 [0.857, 1.062] | 0.319 | 5.68% [-1.21%, 13.57%] Lower |

For go and no-go, conventional averaging (CA) yielded sRMSE of 0.631, 95%CI [0.554, 0.720] and 0.705, 95%CI [0.627, 0.783], respectively. Distance-weighted averaging systematically increased deviation relative to CA, with deviations ranging from 1.4% to 47.1% depending on condition and sharpness parameter. Channel-by-channel heatmaps (**Figure 9 A2-A4, B2-B4**) showed comparable spatial patterns across averaging schemes, indicating preserved lag structure despite numerical differences in aggregate error.

For go (select), CA produced sRMSE of 1.123, 95%CI [0.939, 1.367]. Distance-weighted averaging reduced deviation systematically: κ=1 achieved 19.2%, 95%CI [8.5%, 29.9%] reduction; κ=2 yielded 4.7%, 95%CI [-9.3%, 16.3%] reduction; κ=3 produced 5.7%, 95%CI [-1.2%, 13.6%] reduction. Heatmaps (**Figure C2-C4**) showed spatial lag patterns similar to the go condition.

**Figure:**
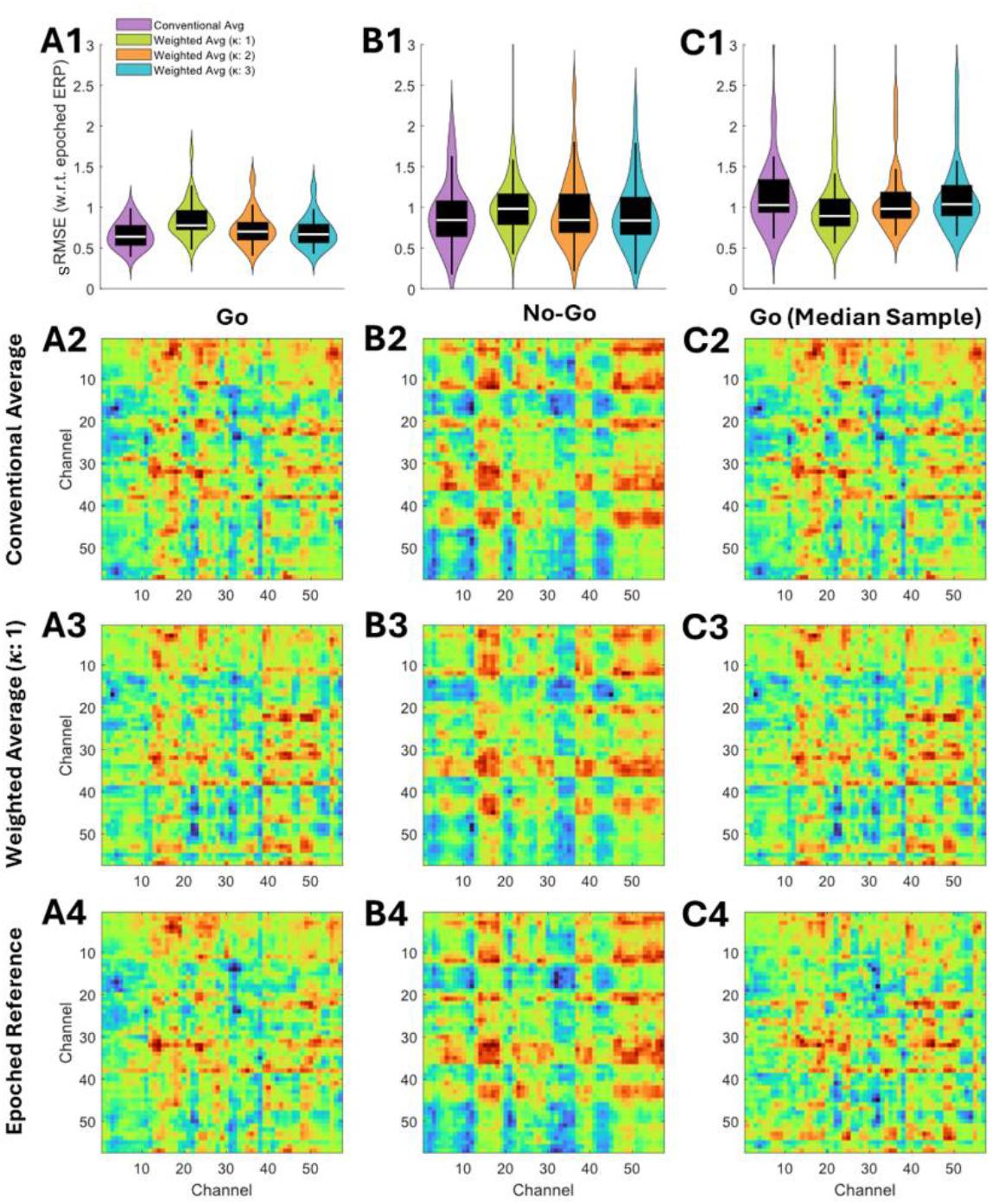
Inter-sensor timing preservation following temporal realignment. Columns indicate trial type: A, go trials; B, no-go trials; C, go trials with restricted cue-to-target intervals ($\pm$100 ms of median). Row 1: sRMSE distributions quantifying lag structure deviation between ERW-aligned and epoched reference cross-covariance matrices. Conventional averaging (purple) and weighted averaging at *κ* ∈ {1, 2, 3} (yellow-green, orange, cyan) show minimal differences. Lower sRMSE indicates better preservation of inter-channel temporal relationships. Rows 2 to 4: Channel-by-channel sRMSE heatmaps for conventional averaging (row 2), weighted averaging *κ* = 1 (row 3), and epoched reference (row 4), averaged across participants. Colour scale indicates magnitude of lag deviation; blue represents minimal perturbation. Heatmap similarity across rows indicates ERW preserves intrinsic lead-lag relationships without introducing spurious phase shifts.

#### 3.2.3 Summary

ERW successfully recovered target-locked event-related potentials from naturalistic data containing substantial trial-to-trial temporal variability, achieving sRMSE values of 0.24 to 0.51 across conditions despite variable cue-to-target intervals spanning 1.5 to 4.1 s. Extended time-course analysis confirmed that temporal realignment sharpened target-locked components without distorting cue-related activity, demonstrating selective correction of jittered events whilst preserving genuine temporal structure. Cross-covariance analysis revealed that ERW preserved intrinsic inter-channel lag relationships following temporal realignment, with channel-by-channel heatmaps showing consistent spatial patterns between ERW-aligned and epoched reference data (sRMSE 0.66 to 1.25), validating ERW’s suitability for subsequent connectivity and causal analyses. Distance-weighted averaging effects were reference-dependent: weighting increased error for go and no-go trials (which incorporated full temporal variability) but decreased error for go (select) trials (derived from temporally restricted subsets), reflecting that weighted averaging naturally emphasised temporally consistent trials rather than indicating inherent method limitations.

## 4 Discussion

ERW successfully recovered temporally jittered responses across simulation and empirical validations. Synthetic data with known ground truth yielded standardised root-mean-square errors of 0.27–0.38 across diverse variability patterns. Distance-weighted averaging provided 5–13% improvements when temporal variability exceeded 100 ms, with optimal sharpness parameter *κ* = 1. Empirical validation using naturalistic go/no-go data with cue-to-target intervals spanning 1.5–4.1 seconds confirmed successful recovery of sequential event-related potentials whilst preserving inter-channel timing relationships.

ERW’s uniqueness lies in operating on latent event representations rather than observed signals. Whilst DTW, Woody filter, and diffeomorphic methods warp neural data directly, and deconvolution approaches model overlap without realignment, ERW aligns template functions encoding experimental design structure. This design-level warping reduces noise susceptibility, enables uniform application across channels to preserve causal relationships, and reconstructs unified sequential responses through conventional averaging, contrasting with decomposition and deconvolution methods that yield separate component or event-type estimates.

ERW is particularly suited to multi-event sequences with variable inter-event timing where preserving inter-channel relationships is critical. By estimating a single warp trajectory per trial applied uniformly across all channels, ERW maintains causal structure essential for connectivity analysis, Granger causality, phase synchrony, and dynamic causal modelling. Channel-specific alignment would violate these causal constraints, rendering subsequent network analyses invalid.

The same warping trajectories can be applied to simultaneously recorded modalities (EEG, MEG, fMRI, pupillometry, skin conductance) provided they share a common event structure. This uniform application distinguishes ERW from signal-level warping methods that would require independent alignment per modality, potentially disrupting cross-modal synchrony.

### 4.1 Runtime

Typical single-subject alignment (64 channels, 60 trials, 1400 time points, K = 8, S = 18) requires approximately 2 minutes on standard workstations. Computational demands increase linearly with trial count and sampling rate, making ERW feasible for typical EEG/MEG, as the procedure does not operate at the level of data; we envisage no need for optimisation for high-density recordings.

### 4.2 Future Directions

Several methodological extensions warrant investigation. The current ERW implementation achieves smooth, monotonic temporal warping but does not strictly satisfy diffeomorphic criteria, which require that both the warping function and its inverse remain smooth everywhere. The loss of diffeomorphism occurs at the step where we enforce monotonicity: to guarantee time always flows forward, we constrain the warping increments to be strictly positive, and the method used to enforce this constraint introduces potential non-differentiable points in the warping function. While this design choice ensures computational efficiency and robust alignment, it sacrifices the bidirectional smoothness required for true diffeomorphism. For standard event-related potential analysis, where primary interest lies in forward-time signal alignment and averaging, the approximate diffeomorphism achieved through DST-parameterised monotonic functions may suffice.

Diffeomorphic extensions could prove advantageous if the problem space were expanded to include uncertainty in the event structure itself. For example, within go/no-go paradigms, response-locked events present an asymmetry: go trials contain clear button-press times, whereas no-go trials lack an overt behavioural marker at corresponding latencies. This complicates direct comparison of response-locked components across conditions. A Bayesian inversion scheme incorporating diffeomorphic warps could simultaneously estimate latent event times and alignment parameters within a unified generative model. By treating event onset times as hidden variables with prior distributions, such a framework would enable probabilistic inference about when covert processing events occur in no-go trials, informed by the temporal structure recovered from go trials. The smooth invertibility guaranteed by diffeomorphic warps would ensure that uncertainty about event timing propagates consistently through the model, maintaining well-defined probability densities under coordinate transformations. This would necessitate integrating observed data directly into the estimation process, moving beyond the current template-matching paradigm to a fully generative model of trial-level temporal variability.

More broadly, Bayesian formulations offer principled handling of alignment uncertainty. Joint estimation of templates and warps within a single hierarchical model would account for inter-trial covariance and uncertainty in event timings rather than treating templates as fixed targets. Bayesian parameter estimation could automatically select DST order K, the number of interpolation steps S, and weighting sharpness κ based on data characteristics through model comparison, eliminating manual tuning.

### 4.3 Conclusion

ERW addresses temporal jitter in event-related response analysis through a simple but conceptually novel approach: warping latent event structures rather than observed neural signals. By aligning template functions encoding experimental design, ERW reduces noise susceptibility, preserves inter-channel causal relationships through uniform trajectory application, and reconstructs unified sequential responses compatible with conventional ERP interpretation. Validation using synthetic data with known ground truth and naturalistic experimental recordings establishes ERW’s efficacy for paradigms involving sequential events with substantial trial-to-trial variability. The method is particularly valuable for multichannel and multimodal recordings where preserving temporal relationships is essential for connectivity and network analyses. ERW provides a principled framework for recovering temporally coherent responses from ecologically valid experimental designs that challenge conventional averaging assumptions.

